# Lassa Virus in the Host Rodent *Mastomys Natalensis* within Urban Areas of N’zerekore, Guinea

**DOI:** 10.1101/616466

**Authors:** LS Karan, MT Makenov, MG Korneev, N Sacko, S Boumbaly, RB Bayandin, AV Gladysheva, K Kourouma, AH Toure, MYu Kartashov, AV Shipovalov, AM Porshakov, M Koulibaly, MY Boiro

## Abstract

Lassa virus is the causative agent of a dangerous zoonotic disease distributed in West Africa. A primary reservoir host of Lassa virus is *Mastomys natalensis*. These mice associate closely with humans and are commonly found in villages. Consequently, previous studies of Lassa virus have focused on rural areas. The prevalence of the virus in large cities has not been studied.

We conducted a study in N’Zerekore city, which has a population of approximately 300,000 residents. Small mammals were captured during a pilot study in May, and the main study was performed in August 2018. Based on the pilot study, we designed and implemented a stratified random sample to investigate the prevalence of Lassa virus among *M. natalensis* in N’Zerekore. The total sampling efforts consisted of 45 and 985 trapping nights in May and August, respectively. Samples of rodent tissues were screened for Lassa virus by RT-PCR.

In May, we trapped 20 rodents, including 19 *M. natalensis*. Viral RNA was detected in 18 *M. natalensis*. In August, 149 small mammals were captured, including 43 *M. natalensis*. The prevalence of Lassa virus among *M. natalensis* in N’Zerekore was 23.3% (CI 95%: 11.8–38.6%). Sequencing showed that the isolates belonged to lineage IV. We detected four Lassa virus hotspots located in different parts of the city. The largest Lassa virus hotspot was found in the neighborhood of the central market, which suggests that the virus was originally introduced into the city through the market.

## Introduction

Lassa virus is the causative agent of Lassa fever, a dangerous zoonotic disease distributed in West Africa. Outbreaks of Lassa fever have emerged in Sierra Leone from 1971–83 and in 1997; Liberia in 1972, 1977, and 1982; Guinea in 1982–83; and Nigeria in 1970, 1974–77, 1989, 1993, 2004 (Fichet-Calvet and Rogers, 2009). In 2018, a new Lassa fever outbreak was reported in Nigeria; as of December 31, 2018, 633 cases were confirmed with 177 deaths (*2018 Lassa fever outbreak in Nigeria, 2018*).

A primary reservoir host of Lassa virus is the multimammate mouse *Mastomys natalensis* (Lecompte et al., 2006; Monath et al., 1974); the virus has been additionally detected in *M. erythroleucus* (Mccormick et al., 1987; Olayemi et al., 2016) and *Hylomyscus pamfi* (Olayemi et al., 2016). Rodents exhibit persistent, asymptomatic infection and shed the virus through feces, urine and saliva. Currently, six lineages of Lassa virus are described: lineages I, II and III circulate in Nigeria, lineage IV was identified in Guinea, Sierra Leone, and Liberia, lineage V (a subclade of lineage IV) circulates in Mali and Côte d’Ivoire, and lineage VI was isolated from *H. pamfi* in Nigeria (Andersen et al., 2015; Bowen et al., 2000; Olayemi et al., 2016); a new variant isolated in Togo is related to lineages II and I/VI in different segments of the genome (Whitmer et al., 2018).

In Guinea, Lassa virus has been found both in rodents (Demby et al., 2001; Fichet-Calvet et al., 2016, 2007; Lecompte et al., 2006) and humans (Bausch et al., 2001). In addition, two studies of Lassa virus seroprevalence were conducted among the Guinean population (Bausch et al., 2001; Lukashevich et al., 1993). Most of these works focused on rural areas, and large cities were not studied. The exception is the work of Demby et al. (Demby et al., 2001), wherein the authors trapped rodents in the town of Kindia (which had a population of approximately 60,000 at the time). The results of their work showed the absence of Lassa virus in rodents in Kindia (Demby et al., 2001).

Therefore, our knowledge of Lassa virus ecology, epidemiology and distribution in West Africa is poorly understood (Gibb et al., 2017); in particular, the role of cities in Lassa fever epidemiology is still to be elucidated. In this work, we studied the prevalence of Lassa virus among rodent hosts within a large city in Guinea.

## Methods

### Small-mammal trapping

The study was conducted in N’Zerekore, Guinea. N’Zerekore has a population of approximately 300,000 and is the second largest city in Guinea. The city is the capital of the N’Zerekore Prefecture and is located in the Guinée forestière region.

We carried out field research in two stages: the pilot study was performed on 5–7 May 2018 (early rainy season), and the main study was performed on 21–31 August 2018 (late rainy season). Small mammals were captured using live traps (Sherman LFA Live Traps, HB Sherman Traps Inc., Tallahassee, FL, USA). During the pilot study, traps were set up for three consecutive nights in three different households in the Tielepolou District of N’Zerekore. The total capture effort in the pilot stage comprised 45 trapping nights.

Based on the pilot study, we designed a stratified random sample to investigate the prevalence of Lassa virus among *M. natalensis* in N’Zerekore. It was shown that in Guinean villages, the distances between recaptures never exceeded 100 m for *M. natalensis* (Mariën et al., 2018). We took this parameter into account and divided the urban territory into 100 x 100 meter squares that were considered sample units. Then, we divided the territory of the city into 4 strata (Fig. 1) as follows:

1. Periphery: The residential area adjacent to undeveloped territories and cultivations that surround the city. All squares that were located no more than 200 meters from the edge of the city were classified as Periphery.
2. Residential area: The residential area (excluding the Periphery) represented mostly by single-story buildings.
3. Marketplaces: The territory of street markets with a large number of small buildings and warehouses. These territories were actively used in the daytime and were empty at night.
4. Cultivation area: Mostly floodplains containing vegetable gardens with rice, corn, manioc, peanuts, etc. Using QGIS software, we implemented this stratification to N’Zerekore and defined the number of sample units in each stratum and in the whole city (Fig. 1), which allowed us to build the stratified sample with proportional allocations. For Marketplaces, we increased the sample fraction due to the small size of this stratum. As a result, the total sample consisted of 123 squares; in each sampled square, we set 5–12 live traps for one night, and the total capture effort comprised 985 trapping nights (Table 1).

**Fig. 1.**
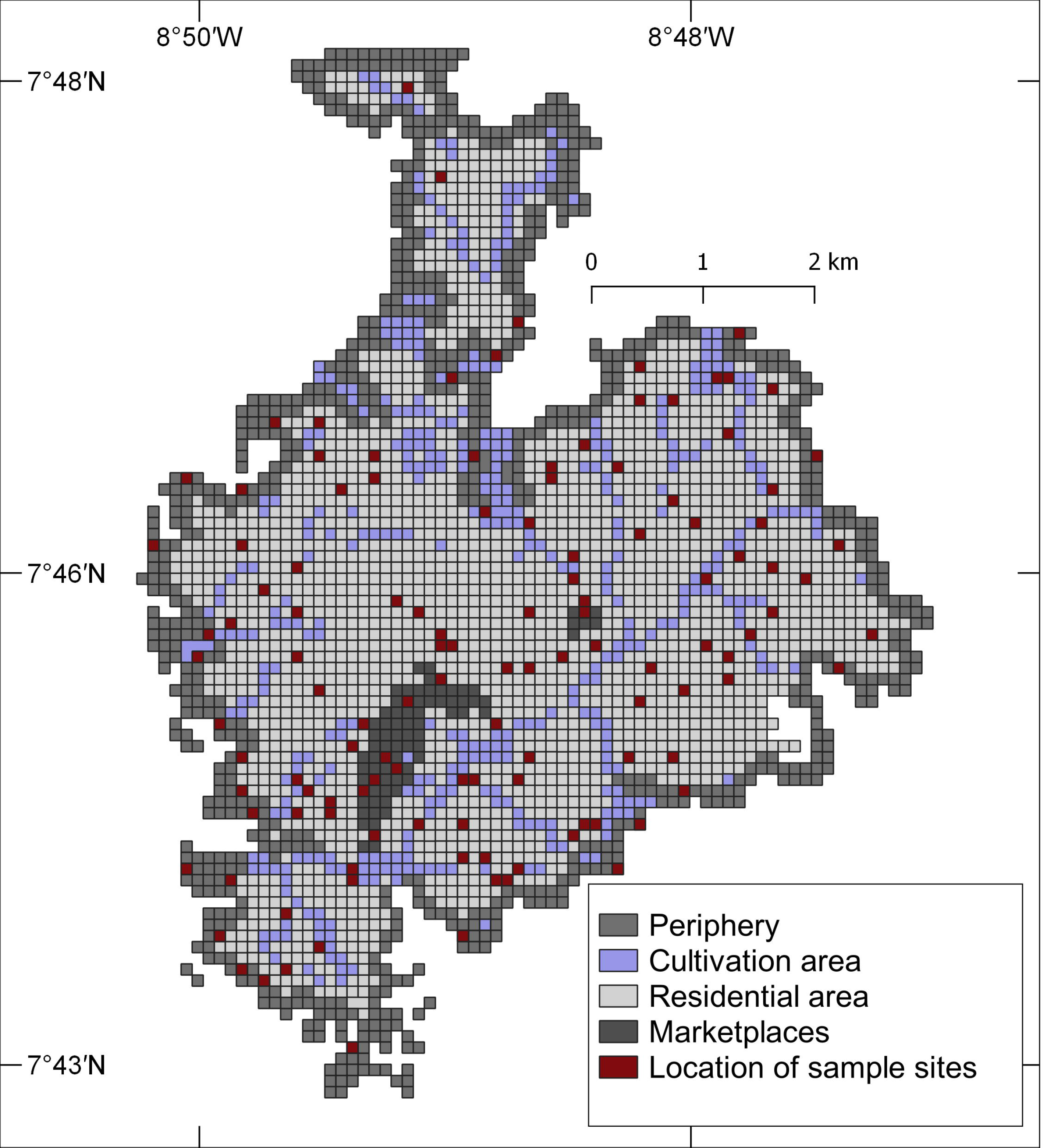
Stratification map of N’Zerekore and sampling site allocations

The main study was conducted on 21–31 August 2018. Standard methods were followed for the safe handling and sampling of small mammals potentially infected with infectious pathogens (Mills et al., 1999).

**Table 1.** The stratified sample for rodent trapping in N’Zerekore

| Stratum | The number of squares (sample units) |  | The number of traps |
| --- | --- | --- | --- |
|  | In the city | In the sample |  |
| Periphery | 786 | 26 | 197 |
| Residential area | 1953 | 76 | 630 |
| Marketplaces | 80 | 7 | 51 |
| Cultivation area | 421 | 14 | 107 |
| Total | 3240 | 123 | 985 |

All trapped small mammals were described morphologically, weighed, and measured (length of head, body, tail, hind foot, and ear) and then typed by sequencing the COI gene from liver autopsies using the following primers: ST–COI–F2 5’–TCYACYAAWCAYAAAGACATTGGAAC–3’ (Makenov et al., 2018) and jgHCO2198 5’–TAIACYTCIGGRTGICCRAARAAYCA–3’ (Geller et al., 2013).

### Sample collection

Blood was collected from the trapped animals in sterile tubes with 0.5 M EDTA by cardiac puncture. Animals were then euthanized, and sections of the liver, spleen, kidney, and lung were obtained through sterile necropsy. Blood samples were centrifuged to separate the plasma and cells, and all blood fractions were stored at 10°C for five days and at minus 28°C for the next 10 days before analysis. Tissue samples were collected in RNAlater (Thermo Fisher Scientific, Waltham, MA, USA) and stored at 6–10°C for 12 days before analysis.

### RNA extraction, polymerase chain reaction and sequencing

We extracted RNA from (1) 100 µl of blood; (2) 10% suspension of brain; (3) and 10% suspension pools of internal organs (liver, spleen, lung and kidney) using a commercial kit (AmpliSens RiboPrep Kit, Central Research Institute of Epidemiology, Moscow, Russia) following the manufacturer’s instructions. Samples were screened for Lassa virus by qRT-PCR using specific primers and probes (Table 2).

**Table 2.**
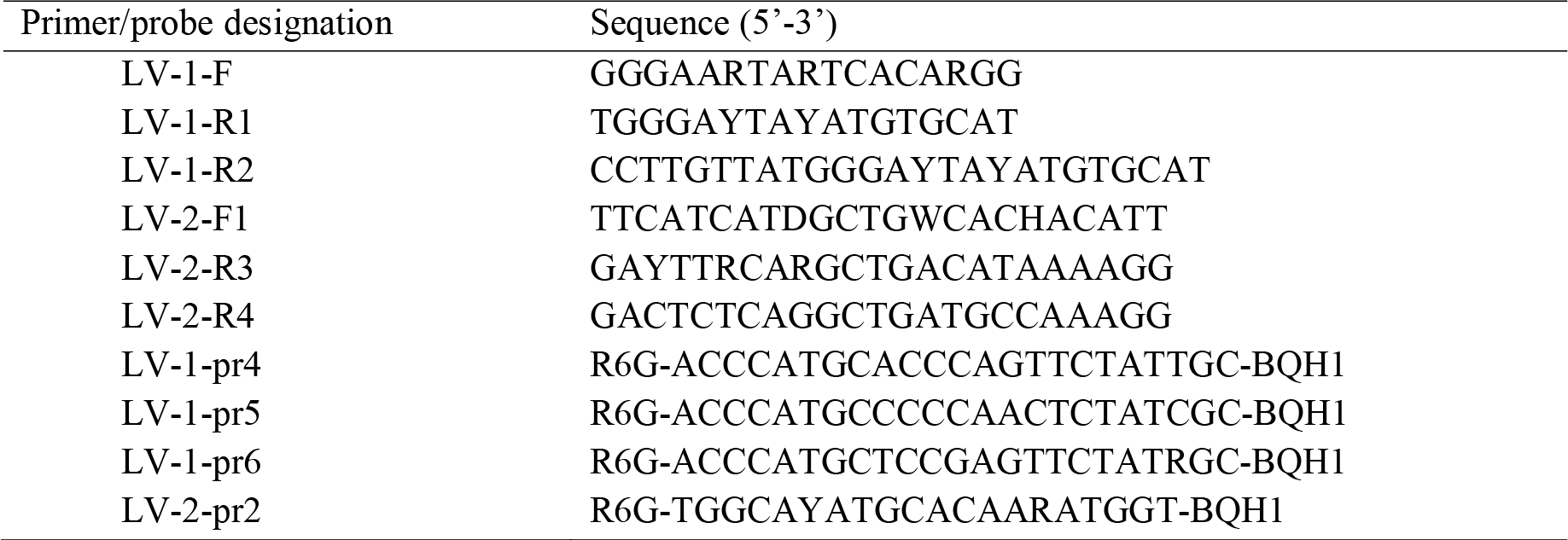
Oligonucleotide primers and probes for duo-target L segment qRT-PCR assay used in the study

From Lassa virus–positive samples, a fragment of the NP coding region was amplified using an RT-PCR protocol (Fichet-Calvet et al., 2016). Purified PCR products were determined be Sanger sequencing. Sequences obtained were deposited in NCBI GenBank under the following accession numbers: MH748572 (M. natalensis COI gene) and MH732623–MH732640 (Lassa virus nucleoprotein gene, S–segment).

### Data analyses

We created a random stratified sample in R using the package ‘survey’ (Lumley, 2004). We estimated the prevalence and confidence intervals using exact Clopper–Pearson methods in the package ‘PropCIs’ (Scherer, 2014). Sequence analysis and multiple alignments were performed using Geneious (version 9.1.7; https://www.geneious.com/). Bayesian coalescent phylogenetic analysis was implemented using MrBayes with substitution model GTR.

## Results

### Small mammals

During the pilot study in May, we trapped two species of rodents, including 19 *M. natalensis* and one *Mus musculus*. Of the 19 *M. natalensis*, 18 yielded positive qRT-PCR signals for Lassa virus. Viral RNA was detected in blood as well as in tissue samples, with the lowest Ct in brain tissue samples (Fig. 2). The concentration of the virus was significantly higher in the brain tissues than in the parenchymal organs (paired t–test: t = −4.99; df = 15; p < 0.001) and in blood (paired t–test: t = − 9.13; df = 13; p < 0.001).

**Fig. 2.**
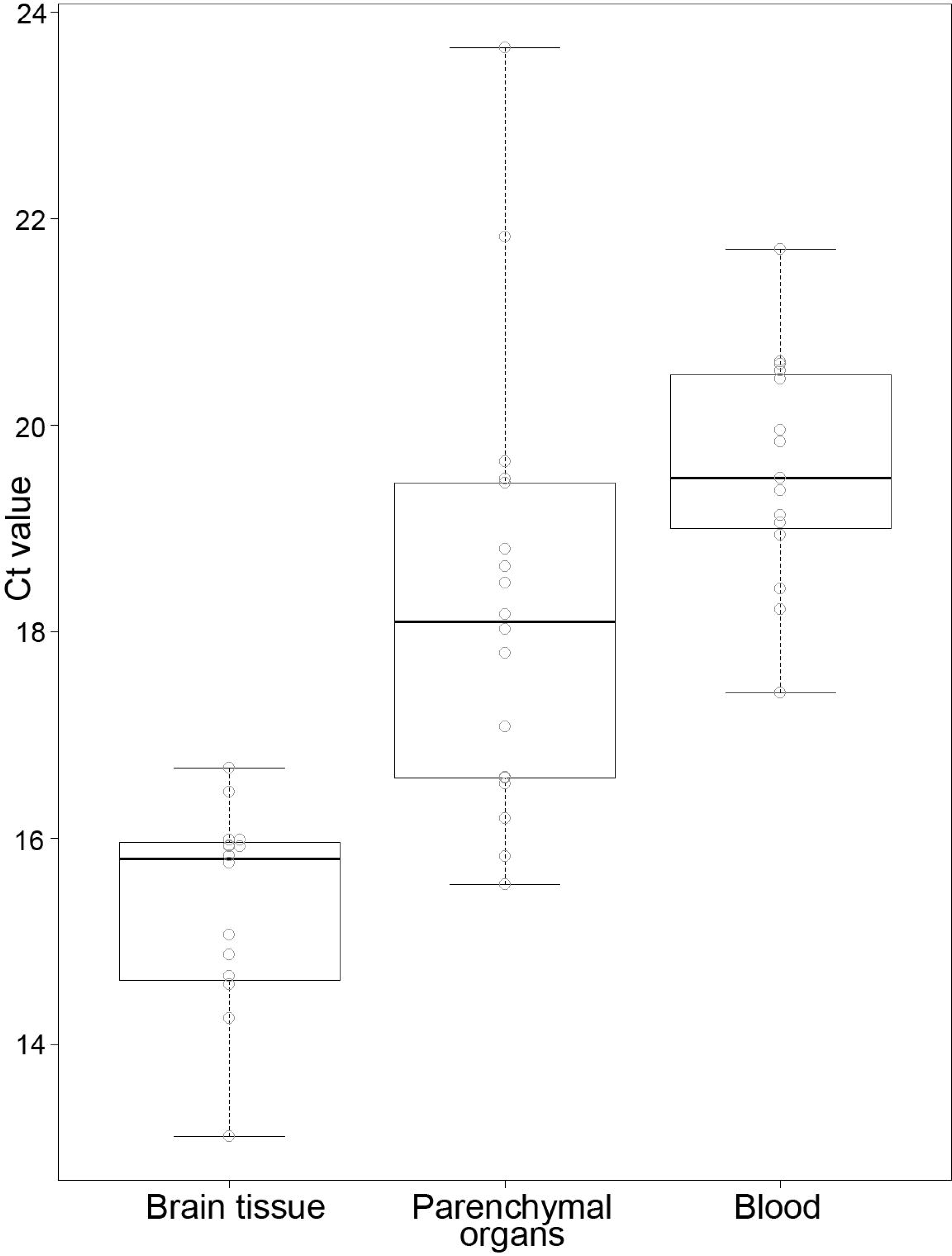
The Ct value of Lassa virus RNA in blood and tissue samples. The solid line represents the median, the box shows the interquartile range (IQR), and the whisker represents 1.5×IQR.

During the main study in August, a total of 149 small mammals representing at least four species were live-trapped (Table 3). Species identification was confirmed by sequencing the COI gene for all *M. natalensis*. For the remaining species, only a few specimens were sequenced. Since the main goal in this work was to study the prevalence of Lassa virus among *M. natalensis*, we did not identify shrews to the species level; they were combined at the genus level as «*Crocidura* spp.». The highest prevalence of *M. natalensis* was in the Residential area, and *R. rattus* and *M. musculus* were most prevalent in Marketplaces (Table 3). In some squares and even in some households, we trapped these three species rodents simultaneously. The blood and internal organs samples from all 43 *M. natalensis* were screened for Lassa virus by qRT-PCR, and 10 samples, which accounted for 23.3% (CI 95%: 11.8–38.6%) of the population, yielded positive signals.

**Table 3.** Number of trapped small mammals by species and by stratum

| Stratum | Number of trapping nights | The number of trapped small mammals / Trapping success* |  |  |  | Total |
| --- | --- | --- | --- | --- | --- | --- |
|  |  | *Mastomys natalensis* | *Rattus rattus* | *Mus musculus* | *Crocidura* spp. |  |
| Periphery | 197 | <u>6</u><br>3.1 | <u>1</u><br>0.5 | <u>7</u><br>3.6 | <u>6</u><br>3.1 | <u>20</u><br>10.2 |
| Residential area | 630 | <u>34</u><br>5.4 | <u>9</u><br>1.4 | <u>36</u><br>5.7 | <u>17</u><br>2.7 | <u>96</u><br>15.2 |
| Marketplaces | 51 | <u>1</u><br>2.0 | <u>5</u><br>9.8 | <u>4</u><br>7.8 | <u>0</u><br>0.0 | <u>10</u><br>1.6 |
| Cultivation area | 107 | <u>2</u><br>1.9 | <u>2</u><br>1.9 | <u>0</u><br>0.0 | <u>19</u><br>17.8 | <u>23</u><br>21.5 |
| Total | 985 | <u>43</u><br>4.4 | <u>17</u><br>1.7 | <u>47</u><br>4.3 | <u>42</u><br>4.8 | <u>149</u><br>15.1 |

\* – Trapping success was estimated as ( $\Sigma$ number of trapped small mammals / $\Sigma$ trapping nights) × 100

Mapping showed that the Lassa virus–positive multimammate mice were trapped in different parts of the city, including in Tielepolou District, where Lassa virus was detected in May (Square–108, Fig. 3). Due to the lack of data, we could not conduct a hotspot analysis and or estimate Getis–Ord *G*_*i*_*. However, visual analysis of the map allowed us to highlight four hotspots (Fig. 3). The largest cluster covered residential neighborhoods directly adjacent to the central market (Grand Marche De N’Zerekore) from the west. This cluster included five squares, in which seven Lassa virus–positive rodents were caught (Fig. 3: Squares 75, 77, 79, 83, 87). One of these squares was located directly within the Grand Marche De N’Zerekore, which is the largest market in the city with high people traffic (Square 87, Fig. 3). The maximum distance between squares with Lassa virus–positive rodents in this cluster was approximately 900 m. The second hotspot was located in the southeastern part of the city, 1.7 km east of the Grand Marche De N’Zerekore (Square 108, Fig. 3). At this site, 18 Lassa virus–positive rodents were caught in May, and one infected specimen was trapped in August. Two additional hotspots were located in the northwestern and northeastern parts of the city, each of which included only one Lassa virus–positive *M. natalensis*.

**Figure 3.**
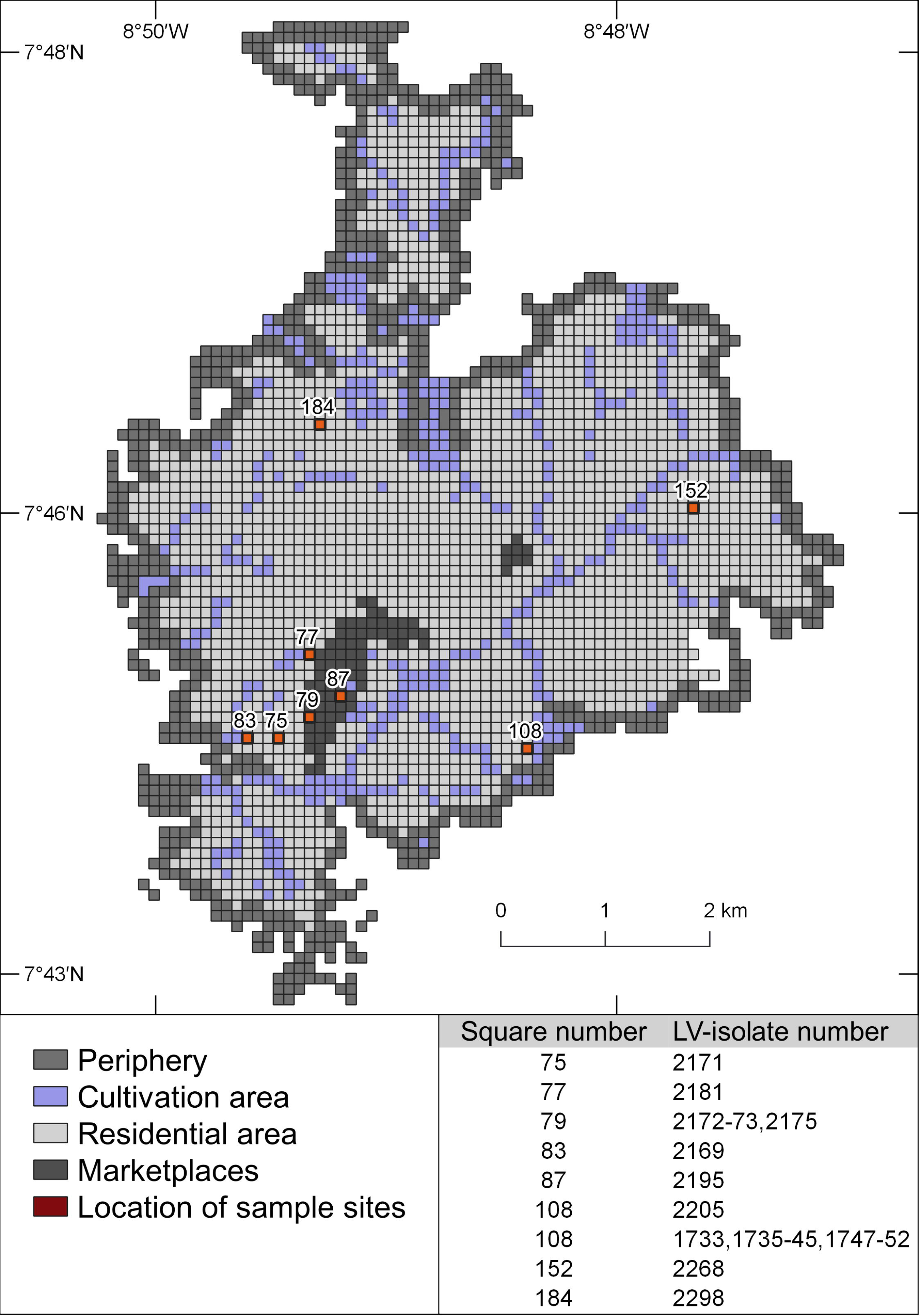
Allocation of Lassa virus–positive *Mastomys natalensis* trapping sites

### Virus characteristics

We genetically characterized the detected Lassa virus by determining its partial NP coding nucleotide sequence (756-bp fragments) and comparing it with other Lassa virus sequences. The obtained viruses belonged to lineage IV; furthermore, all obtained isolates formed a separate phylogenetic branch (Fig. 4). Genetic diversity between the N’Zerekore strains in 2018 approached 5% nucleotide divergence. The lowest diversity (0.7%) was found between isolates from Tielepolou (Square 108, Fig. 3), and the highest diversity (4.7%) was recorded between isolates obtained from neighborhoods of the Grand Marche De N’Zerekore (Fig. 3: Squares 75, 77, 79, 83, 87). For comparison, the genetic distances between N’Zerekore isolates and other Guinean Lassa virus strains including Macenta (10.2–11.4%), Кissidougou (11.7–13.2%), Madina-Oula (11.4–14.0%), and Faranah (11.2–16.0%) were significantly higher.

**Figure 4.**
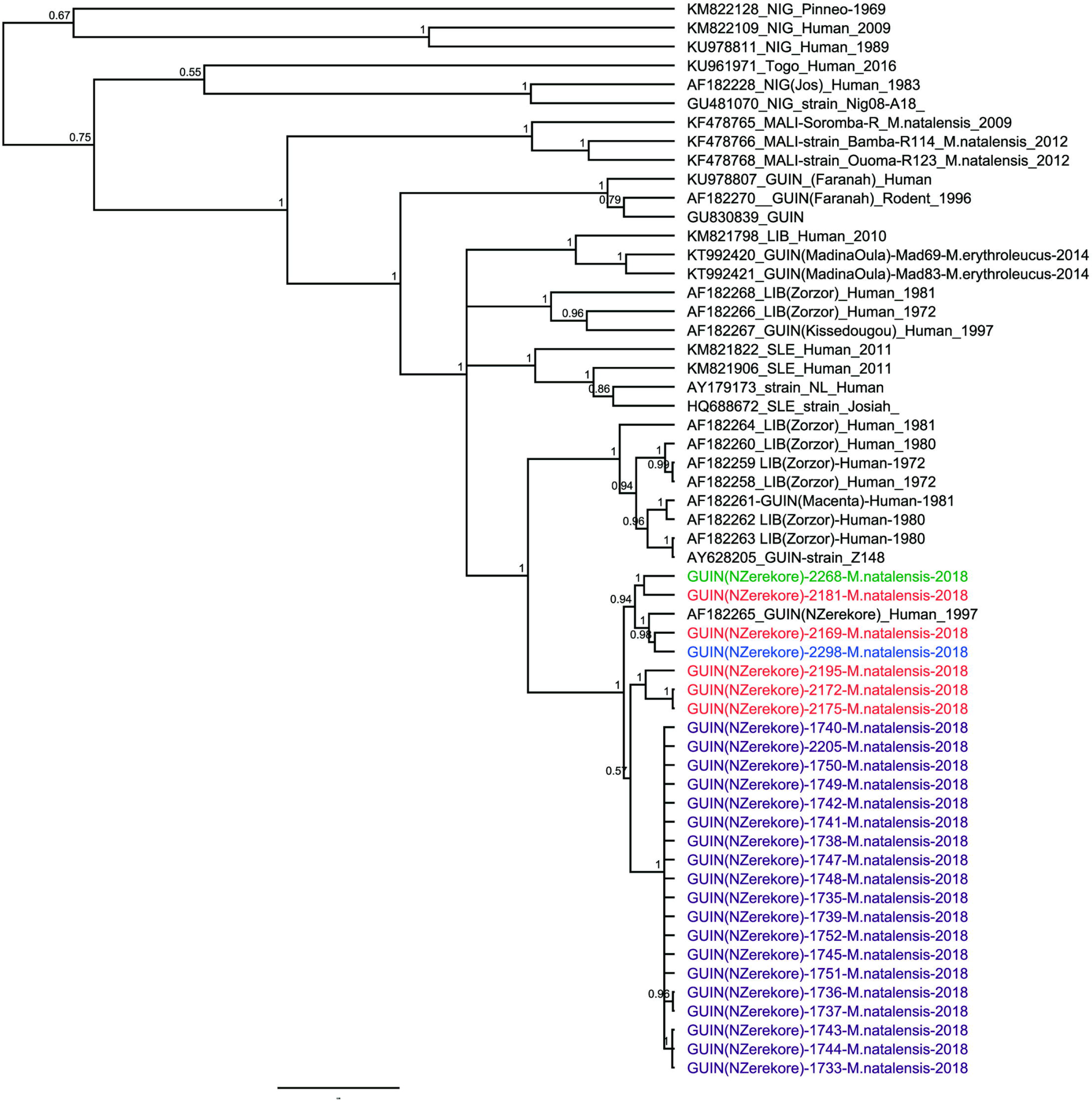
Phylogenetic analysis of Lassa virus NP sequences (756 bp) from 26 *Mastomys natalensis* studied in this work. The tree was inferred by using Bayesian Markov Chain Monte Carlo method, with GTR+gamma model. Scale bar indicates 2% divergence. The colors represent different hotspots: Purple – Tilépoulou District, Red – The central market, Blue and Green – on the northwestern and northeastern part of the city, respectively.

## Discussion

We detected a very high prevalence of Lassa virus in *M. natalensis* in the Tielepolou District of N’Zerekore in the early rainy season (18/19 of trapped rodents). Further study showed that the prevalence of Lassa virus in *M. natalensis* in the city was 23.3% (CI 95%: 11.8–38.6%). Therefore, we have the first reliable evidence of the presence of Lassa virus–infected rodents in a large city in West Africa. In previous works, Lassa virus was detected in *M. natalensis* in rural areas in villages of Guinea (Demby et al., 2001; Fichet-Calvet et al., 2007; Lecompte et al., 2006), Sierra Leone (Bowen et al., 2000; Leski et al., 2015; Mccormick et al., 1987; Monath et al., 1974; Olayemi et al., 2016), Nigeria (Agbonlahor et al., 2017; Olayemi et al., 2016), Côte d’Ivoire (Kouadio et al., 2015), and Mali (Safronetz et al., 2013); to date, the pathogen has not been found in rodents in urban areas.

The presence of Lassa virus-infected rodents in urban areas with a high human population density could lead to a new Lassa fever outbreak in Guinea. In Guinea, only one Lassa fever outbreak has been described from the subprefecture Madina-Oula in 1982–83 (Boiro et al., 1987; Sochinsky et al., 1983). The outbreak involved at least 11 villages near the border with Sierra Leone; 360 inhabitants were affected, and 138 of them died (Boiro et al., 1987). However, it is worth mentioning that the causative agent of this outbreak was not determined. Antibodies to Lassa virus were detected in three convalescent patients (*n* = 22), while convalescent sera from 15 of 79 patients (19%) were positive for antibodies to ebolaviruses (Boiro et al., 1987; Sochinsky et al., 1983). A high level of seroprevalence to Lassa virus (34.5%) was detected in this region eight years after the outbreak (Lukashevich et al., 1993). These data support the hypothesis of a Lassa fever outbreak in Madina Oula in 1982–83, but the etiology of this outbreak is unclear. Subprefecture Madina Oula is far from N’Zerekore (at a distance of 450 km). In the Guinée forestière region and the N’Zerekore Prefecture, a Lassa fever outbreak has not been registered thus far.

A serosurvey for Lassa virus–specific IgG antibodies in Guinea showed that in 1990–92, a high frequency of IgG–positive samples was found in the Guinée forestière region (mean prevalence was 33%) and in the Faranah Prefecture (mean prevalence was 36%) (Lukashevich et al., 1993). Kerneis et al. (Kernéis et al., 2009) studied Lassa virus seroprevalence in the Guinée forestière region in 2000. The authors divided the studied settlements into rural and urban areas, the latter of which was defined as a settlement with a population size greater than 3,000 and easy road access (Kernéis et al., 2009). The results showed that Lassa virus seroprevalence in rural and urban areas, at 13% and 10%, respectively, was approximately the same (Kernéis et al., 2009). Despite the fact that a Lassa fever outbreak has not been recorded in Guinea since 1983, all of the evidence suggests contact of the local population with the virus. In fact, 22 confirmed cases of acute Lassa fever in Guinea were reported in 1996–1999, the majority of which were found in the Guinée forestière region, with 11 cases presenting at the hospital of N’Zerekore and six cases at the hospital of Kissidougou (Bausch et al., 2001). Lassa virus–positive rodents (*M. natalensis* and/or *erythroleucus*) have been captured in the Faranah and N’Zerekore regions and in the Madina-Oula subprefecture (Demby et al., 2001; Fichet-Calvet et al., 2007; Lecompte et al., 2006; Olayemi et al., 2016).

In previous works, it has been shown that the Lassa virus from Guinea belongs to lineage IV (Bowen et al., 2000). Bowen et al. (Bowen et al., 2000) also found that the Lassa virus genetic distance within lineage IV correlates with geographical distance. Our isolates of Lassa virus also belonged to lineage IV and had the closest relation with a strain isolated from a patient in N’Zerekore in 1997 (GenBank accession number AF182265). Apparently, the Lassa virus in N’Zerekore forms a separate geographically related branch of the virus.

Mapping the sites where infected rodents were caught allowed us to highlight four Lassa virus hotspots within N’Zerekore. These hotspots were located in different parts of the city and were relatively distant from each other. Furthermore, sequencing showed that the Lassa virus from different hotspots had genetic differences. The hotspot in the southeastern part of the city (Tielepolou District) included 19 isolates of Lassa virus (Square 108, Fig. 3). These isolates had the lowest genetic distances between each other, with four substitutions on the 756 bp fragment of the nucleoprotein gene, and significant distances from isolates from the rest of the hotspots, with approximately 35 substitutions on the 756 bp fragment (Fig. 4). We assume that all of the isolates represent secondary cases produced by one virus introduction event. Since this hotspot was close to the city limits (approximately 250 m), we suggest that the virus was introduced to Tielepolou District from neighboring rural areas.

The hotspot from the west side of the central market (Grand Marche De N’Zerekore) (Fig. 3) was formed by isolates that were closely located but had relatively low genetic homology. The scenario that all isolates originated from one founding virus does not apply for this hotspot because of the high genetic heterogeneity. Apparently, several different founding viruses were introduced to this hotspot at different times. Phylogenetic and spatial analyses allow us to suggest that Lassa virus was introduced into the city through the market. Indeed, the largest Lassa virus hotspot with the highest genetic diversity was found in the neighborhood of the Grand Marche De N’Zerekore (Fig. 3). The market is a place where people store and sell rice, peanuts, palm oil, fish and other products imported daily from the villages. Perhaps, infected rodents or contaminated grain entered the city through transport from the countryside. Hotspots in the northwestern and northeastern parts of the city were far from Tielepolou District and the central market (Fig. 3: Squares 184 and 152, respectively); however, the isolates from these hotspots were genetically close to the virus from the hotspot near the central market. Most likely, independent Lassa virus introduction events from rural areas occurred, or the virus was introduced into the market at first and then to the northern districts of the city. To test this assumption, it is necessary to obtain virus isolates from the villages of the Guinée forestière region and carry out a phylogeographic analysis.

The concentration of viral RNA was significantly higher in the rodents’ brain tissue than in blood and the parenchymal organs. Based on these findings, we recommend using rodents’ brain tissue for viral RNA extraction. This method could increase the sensitivity of the subsequent PCR assay or be helpful for sequencing. In previous works, quantitative assessments of Lassa virus in different host tissues were conducted in experiments using virological techniques (Henderson et al., 1972; Stephenson et al., 1984). In an experiment with Swiss albino mice, the highest virus concentration was in the brain, lung and muscle (Henderson et al., 1972). In an experiment with aerosol–induced Lassa virus infections in outbred guinea pigs, the highest virus concentration was in the lungs, upper respiratory tract, and spleen (Stephenson et al., 1984). In works with wild trapped reservoir rodents, researchers extracted viral RNA from rodent blood (Agbonlahor et al., 2017; Fichet-Calvet et al., 2007; Olayemi et al., 2016), spleen (Fichet-Calvet et al., 2007; Leski et al., 2015) or liver tissues (Lecompte et al., 2006). Brain tissue has not been used thus far for these proposals.

Our results reveal at least two important consequences. First, we need further studies of Lassa virus prevalence in rodents as well as human Lassa virus seroprevalence studies in cities and towns of West Africa, which will help us assess the danger of Lassa virus foci within urban areas and develop an action plan with respect to outbreak prevention. Second, our findings raise the following question: why has a Lassa fever outbreak not already emerged in N’Zerekore? In our study, we detected high concentrations Lassa virus in 28 trapped *M. natalensis* in 2018 within the city limits, but a Lassa fever outbreak in N’Zerekore has not followed. Our findings highlight the importance of further study of the role of cities in Lassa fever epidemiology and the barriers (biological, geographical, ecological) that limit Lassa virus and prevent Lassa fever outbreaks.

## References

1. 2018 Lassa fever outbreak in Nigeria, 2018.

2. Agbonlahor, D.E., Erah, A., Agba, I.M., Oviasogie, F.E., Ehiaghe, A.F., Wankasi, M., Eremwanarue, O.A., Ehiaghe, I.J., Ogbu, E.C., Iyen, R.I., Abbey, S., Tatfeng, M.Y., Uhunmwangho, J., 2017. Prevalence of lassa virus among rodents trapped in three south-south states of Nigeria. J. Vector Borne Dis. 54, 146–150.

3. Andersen, K.G., Shapiro, B.J., Matranga, C.B., Sealfon, R., Lin, A.E., Moses, L.M., Folarin, O.A., Goba, A., Odia, I., Ehiane, P.E., Momoh, M., England, E.M., Winnicki, S., Branco, L.M., Gire, S.K., Phelan, E., Tariyal, R., 2015. Clinical sequencing uncovers origins and evolution of Lassa virus. Cell 162, 738–750. https://doi.org/10.1016/j.cell.2015.07.020.Clinical

4. Bausch, D.G., Demby, A.H., Coulibaly, M., Kanu, J., Goba, A., Bah, A., Condé, N., Wurtzel, H.L., Cavallaro, K.F., Lloyd, E., Baldet, F.B., Cissé, S.D., Fofona, D., Savané, I.K., Tolno, R.T., Mahy, B., Wagoner, K.D., Ksiazek, T.G., Peters, C.J., Rollin, P.E., 2001. Lassa Fever in Guinea: I. Epidemiology of Human Disease and Clinical Observations. Vector borne zoonotic Dis. 1, 269–281.

5. Boiro, I., Lomonossov, N., Sotsinski, V., Constantinov, O., Tkachenko, E., Inapogui, A., Balde, C., 1987. Elements de recherches clinico-epidemiologiques et de laboratoire sur les fievres hemorragiques en Guinee. Bull. La Soc. Pathol. Exot. 80, 607–612.

6. Bowen, M.D., Rollin, P.E., Ksiazek, T.G., Hustad, H.L., Bausch, D.G., Demby, A.H., Bajani, M.D., Peters, C.J., Nichol, S.T., 2000. Genetic Diversity among Lassa Virus Strains. J. Virol. 74, 6992–7004.

7. Demby, A.H., Inapogui, A., Kargbo, K., Koninga, J., Kourouma, K., Kanu, J., Coulibaly, M., Wagoner, K.D., Ksiazek, T.G., Peters, C.J., Rollin, P.E., Bausch, D.G., 2001. Lassa fever in Guinea: II. Distribution and prevalence of Lassa virus infection in small mammals. Vector borne zoonotic Dis. 1, 283–297. https://doi.org/10.1080/15476286.2016.1256535

8. Fichet-Calvet, E., Lecompte, E., Koivogui, L., Soropogui, B., Doré, A., Kourouma, F., Sylla, O., Daffis, S., Koulémou, K., Meulen, J. Ter, 2007. Fluctuation of Abundance and Lassa Virus Prevalence in *Mastomys natalensis* in Guinea, West Africa. Vector-Borne Zoonotic Dis. 7, 119–128. https://doi.org/10.1089/vbz.2006.0520

9. Fichet-Calvet, E., Ölschläger, S., Strecker, T., Koivogui, L., Becker-Ziaja, B., Camara, A.B., Soropogui, B., Magassouba, N., Günther, S., 2016. Spatial and temporal evolution of Lassa virus in the natural host population in Upper Guinea. Sci. Rep. 6, 1–6. https://doi.org/10.1038/srep21977

10. Fichet-Calvet, E., Rogers, D.J., 2009. Risk maps of lassa fever in West Africa. PLoS Negl. Trop. Dis. 3. https://doi.org/10.1371/journal.pntd.0000388

11. Geller, J., Meyer, C., Parker, M., Hawk, H., 2013. Redesign of PCR primers for mitochondrial cytochrome c oxidase subunit I for marine invertebrates and application in all-taxa biotic surveys. Mol. Ecol. Resour. 13, 851–861. https://doi.org/10.1111/1755-0998.12138

12. Gibb, R., Moses, L.M., Redding, D.W., Jones, K.E., 2017. Understanding the cryptic nature of Lassa fever in West Africa. Pathog. Glob. Health 111, 276–288. https://doi.org/10.1080/20477724.2017.1369643

13. Henderson, B.E., Gary, G.W., Kissling, J.. E., Frame, J.D., Carey, D.E., 1972. LASSA FEVER VIROLOGICAL AND SEROLOGICAL STUDIES. Trans. R. Soc. Trop. Med. Hyg. 66, 409–416.

14. Kernéis, S., Koivogui, L., Magassouba, N., Koulemou, K., Lewis, R., Aplogan, A., Grais, R.F., Guerin, P.J., Fichet-Calvet, E., 2009. Prevalence and risk factors of lassa seropositivity in inhabitants of the Forest Region of Guinea: A cross-sectional study. PLoS Negl. Trop. Dis. 3. https://doi.org/10.1371/journal.pntd.0000548

15. Kouadio, L., Nowak, K., Akoua-Koffi, C., Weiss, S., Allali, B.K., Witkowski, P.T., Krüger, D.H., Couacy-Hymann, E., Leendertz, F.H., Allali, B.K., 2015. Lassa virus in multimammate rats, Côte d’Ivoire, 2013. Emerg. Infect. Dis. 21, 1481–1483. https://doi.org/10.3201/eid2108.150312

16. Lecompte, E., Fichet-Calvet, E., Daffis, S., Koulémou, K., Sylla, O., Kourouma, F., Doré, A., Soropogui, B., Aniskin, V., Allali, B., Kan, S.K., Lalis, A., Koivogui, L., Günther, S., Denys, C., Ter Meulen, J., 2006. Mastomys natalensis and Lassa fever, West Africa. Emerg. Infect. Dis. 12, 1971–1974. https://doi.org/10.3201/eid1212.060812

17. Leski, T.A., Stockelman, M.G., Moses, L.M., Park, M., Stenger, D.A., Ansumana, R., Bausch, D.G., Lin, B., 2015. Sequence variability and geographic distribution of Lassa Virus, Sierra Leone. Emerg. Infect. Dis. 21, 609–618. https://doi.org/10.3201/eid2104.141469

18. Lukashevich, I.S., Clegg, J.C.S., Sidibe, K., 1993. Lassa Virus Activity In Guinea - Distribution Of Human Antiviral Antibody-Defined Using Enzyme-Linked-Immunosorbent-Assay With Recombinant Antigen. J. Med. Virol. 40, 210–217. https://doi.org/10.1002/jmv.1890400308

19. Lumley, T., 2004. Analysis of complex survey samples. J. Stat. Softw. 9, 1–19.

20. Makenov, M.T., Karan, L.S., Shashina, N.I., Akhmetshina, M.B., Zhurenkova, O.B., Kholodilov, I.S., Karganova, G.G., Smirnova, N.S., Grigoreva, Y.E., Yankovskaya, Y.D., Fyodorova, M. V, 2018. First detection of tick-borne encephalitis virus in Ixodes ricinus ticks and their rodent hosts in Moscow, Russia. bioRxiv 480475. https://doi.org/10.1101/480475

21. Mariën, J., Kourouma, F., Magassouba, N., Leirs, H., Fichet-Calvet, E., 2018. Movement Patterns of Small Rodents in Lassa Fever-Endemic Villages in Guinea. Ecohealth 1–12. https://doi.org/10.1007/s10393-018-1331-8

22. Mccormick, J.B., Webb, P.A., Krebs, J.W., Johnson, K.M., Smith, E.S., Africa, W., 1987. A Prospective Study of the Epidemiology and Ecology of Lassa Fever carried out primarily in the eastern province of Sierra. J. Infect. Dis. 155, 437–444.

23. Mills, J.N., Childs, J.E., Ksiazek, T.G., Peters, C.J., Velleca, W.M., 1999. Methods for trapping and sampling small mammals for virologic testing. Prevention 61. https://doi.org/10.1017/CBO9781107415324.004

24. Monath, T.P., Newhouse, V.F., Kemp, G.E., Setzer, H.W., Cacciapu.A, 1974. Lassa Virus Isolation From Mastomys Natalensis Rodents During an Epidemic in Sierra-Leone. Science (80-.). 185, 263–265. https://doi.org/10.1126/science.185.4147.263

25. Olayemi, A., Cadar, D., Magassouba, N., Obadare, A., Kourouma, F., Oyeyiola, A., Fasogbon, S., Igbokwe, J., Rieger, T., Bockholt, S., Jérôme, H., Schmidt-Chanasit, J., Garigliany, M., Lorenzen, S., Igbahenah, F., Fichet, J.N., Ortsega, D., Omilabu, S., Günther, S., Fichet-Calvet, E., 2016. New Hosts of The Lassa Virus. Sci. Rep. 6, 1–6. https://doi.org/10.1038/srep25280

26. Safronetz, D., Sogoba, N., Lopez, J.E., Maiga, O., Dahlstrom, E., Zivcec, M., Feldmann, F., Haddock, E., Fischer, R.J., Anderson, J.M., Munster, V.J., Branco, L., Garry, R., Porcella, S.F., Schwan, T.G., Feldmann, H., 2013. Geographic Distribution and Genetic Characterization of Lassa Virus in Sub-Saharan Mali. PLoS Negl. Trop. Dis. 7, 4–12. https://doi.org/10.1371/journal.pntd.0002582

27. Scherer, R., 2014. PropCIs: Various confidence interval methods for proportions. R package version 0.2-5. [WWW Document]. URL https://cran.r-project.org/package=PropCIs

28. Sochinsky, V.A., Legonkov, Y.A., Konde, K., Butenko, A.M., Kamara, M., Fidarov, F.M., 1983. A clinical-epidemiological study of an acute disease with hemorrhagic syndrome which occured in Madina Oula District, Kindia region (1982) (In Russian), in: Arboviruses, Parasitic Diseases, and Bacterial Infections in Republic of Guinea. Conakry, pp. 65–69.

29. Stephenson, E., Larson, E., JW, 1984. Effect of Environmental Factors on Aerosol-Induced Lassa Virus Infection. J. Med. Virol. 14, 295–303.

30. Whitmer, S.L.M., Strecker, T., Cadar, D., Dienes, H.P., Faber, K., Patel, K., Brown, S.M., Davis, W.G., Klena, J.D., Rollin, P.E., Schmidt-Chanasit, J., Fichet-Calvet, E., Noack, B., Emmerich, P., Rieger, T., Wolff, S., Fehling, S.K., Eickmann, M., Mengel, J.P., Schultze, T., Hain, T., Ampofo, W., Bonney, K., Aryeequaye, J.N.D., Ribner, B., Varkey, J.B., Mehta, A.K., Lyon, G.M., Kann, G., De Leuw, P., Schuettfort, G., Stephan, C., Wieland, U., Fries, J.W.U., Kochanek, M., Kraft, C.S., Wolf, T., Nichol, S.T., Becker, S., Ströher, U., Günther, S., 2018. New lineage of Lassa virus, Togo, 2016. Emerg. Infect. Dis. 24, 599–602. https://doi.org/10.3201/eid2403.171905

